# Song Richness Index: A measure to understand the diversity and repetition of notes in a birdsong

**DOI:** 10.1101/2021.12.18.472638

**Authors:** Suyash Sawant, Chiti Arvind, Viral Joshi, V. V. Robin

**Affiliations:** Department of Biology, Indian Institute of Science Education and Research (IISER) Tirupati, Transit campus Karakambadi Road, Tirupati, 517507, India

**Keywords:** Acoustic Complexity, Birdsongs, Note Diversity, Note Repetitions, Shannon’s Diversity, Simpson’s Diversity

## Abstract

Birdsong plays an important role in mate attraction and territorial defense. Many birds, especially Passerines, produce varying sequences of multiple notes resulting in complex songs. Studying the diversity of notes within these songs can give insights into an individual’s reproductive fitness. We first looked at the previously described and commonly used diversity measures to understand the possible case-specific limitations. We then developed a new diversity measure-Song Richness Index (SRI). We compared SRI with three measures of diversity using all possible combinations of notes to understand the case-specific advantages and limitations of all approaches. Simulating all possible combinations gave us insights into how each diversity measure works in a real scenario. SRI showed an advantage over conventional measures of diversity like Note Diversity Index (NDI), Shannon’s Equitability (SH), and Simpson’s Diversity (SI), especially in the cases where songs are made up of only one type of repetitive note.

## Introduction

Birdsongs are one of the critical factors in understanding the ecology and behavior of an individual. Minor acoustic elements, called notes, give rise to the complex sequences of songs. Studying the diversity of these notes can provide us with insights into the various behaviors, including mate selection (László Zsolt Garamszegi, Zsebok, and Török 2012; Boogert, Giraldeau, and Lefebvre 2008), defending territories (Catchpole 1987; Kroodsma and Byers 1991) and warning against predators (Demartsev et al. 2014). Studies have also shown that birds with more diverse songs have a higher reproductive success (Nowicki, Peters, and Podos 1998; Cramer 2013). But given the dynamic nature of these songs, it has always been a challenge to have one robust method to quantify the note diversity (Kershenbaum and Garland 2015; Mikula, Petrusková, and Albrecht 2018).

Kershenbaum et al. (2016) presented various approaches to measuring birdsong complexity which can be categorized into two major groups-measures of sequential variations of notes (order) and measures of the diversity of notes within a song (Sawant et al., 2021). The diversity of birdsongs is often studied in two ways; a) Eventual note diversity-the complexity across songs bouts and b) Immediate note diversity-the complexity within individual song bout. An individual’s entire repertoire comprises the eventual note diversity (L. Z. Garamszegi, Balsby, and Bell 2005; Zeng et al. 2007; Benedict and Najar 2019; Keen et al. 2021), but it does not doesn’t capture the diversity of notes within each song. For example, it is possibel that a bird with a large number of note types (eventual note diversity), may repeat the same notes mutliple times within a song resulting in a low immediate diversity. The immediate diversity of the notes in birdsongs is often studied using some of the conventional measures-Shannon’s Diversity (Kershenbaum 2014; Sasahara et al. 2015; Silva et al. 2000), Simpson’s Diversity (Simpson 1949; Magnussen and Boyle 1995), and the types of notes within a song or Note Diversity (Spencer et al. 2003; Sawant et al. 2021; L. Z. Garamszegi, Balsby, and Bell 2005; Najar and Benedict 2019).

These measures of immediate diversity have shown promising results in understanding the diversity across multiple fields of biology. But as these were primarily designed and used to quantify the diversity of different data sets (e.g., species diversity in community ecology), using them to understand the acoustic diversity of songs has a few limitations. One major limitation is when songs are made of only one note type, and such a song (varying total notes) is assigned the exact value of zero diversity with these conventional measures. Though only one note is repeated in this case, the number of repetitions can have significance in communication, and all such instances cannot be considered the same.

In this study, we describe a new diversity measure, the Song Richness Index (SRI), that overcomes the issue mentioned above regarding the conventional measures of diversity. SRI works similarly to Note Diversity Index (NDI), Shannon’s Equitability (SH), and Simpson’s Diversity (SH) but shows significant case-specific advantages. We highlight these advantages by comparing the SRI with NDI, SH, and SI using sequences of songs created using all possible combinations of note types. We also elaborate on the limitations of each method.

## Methods

### Song Richness Index (SRI)

For a song with N_C_ number of notes, N_T_ note types, the Song Richness Index (SRI) is defined as:

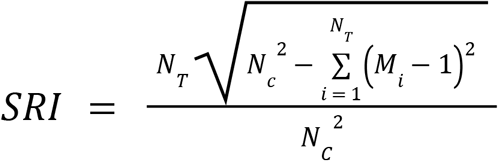

Here, M_i_ represents the frequency of occurrences of each note type, i ∈ (1, N_T_). SRI quantifies the song diversity based on the diversity and repetition of each note type by penalizing each instance of a note type repetition. The numerator which quantifies the note repetition is normalized with N_C_^2^, which results in the output values of SRI ∈ (0,1). For the songs of same length (note count), the higher values of SRI represent high diversity and less repetition, and the lower values represent low diversity and high repetition.

### Comparison of SRI with three conventionally used diversity indices

We created songs made of all possible combinations of notes, ranging from 2 to 10 notes in a song. The notes were represented as distinct elements (A, B, C,…). We then calculated the SRI values for all the possible combinations to understand the case-specific advantages and limitations. We also calculated the diversity of these songs using three widely used measures of diversity: Note Diversity Index (NDI)-the ratio of note types and note count (N_T_ / N_C_), Shannon’s Equitability 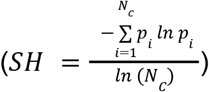 and, Simpson’s Diversity 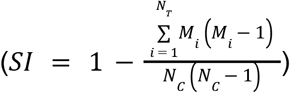 (where 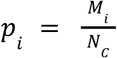 is the stationary probabilities of each note in the song).

We then analyzed the diversity values given by the three measures across different access that include i) songs of different lengths with all combinations of notes, ii) songs of different lengths made of only one note type (repeating), and iii) songs made of same note types and lengths but a varying number of repetitions. We looked at the case-specific limitations of all methods in these scenarios.

## Results

To understand the advantages and limitations of these methods, we compared the Song Richness Index (SRI) with three most commonly used diversity measures-Note Diversity Index (NDI), Shannon’s Equitability (SH), and Simpson’s Diversity (SI) using sequences of notes made of all possible combinations.

For songs made of three, four, and five notes, there are three, five, and seven structural groups of combinations that are possible. When the diversity values for all these songs were calculated, they showed similar diversity patterns with songs with more notes types having higher diversity values. For a few case-specific examples, songs ‘AAAB and AABB’, ‘AAAAB and AAABB’ and ‘AAABC and AABBC’, NDI values were the same suggesting the identical diversity. SH, SI, and SRI could distinguish between these songs based on the number of repetitions of each note type (Fig. 1a).

**Figure 1:**
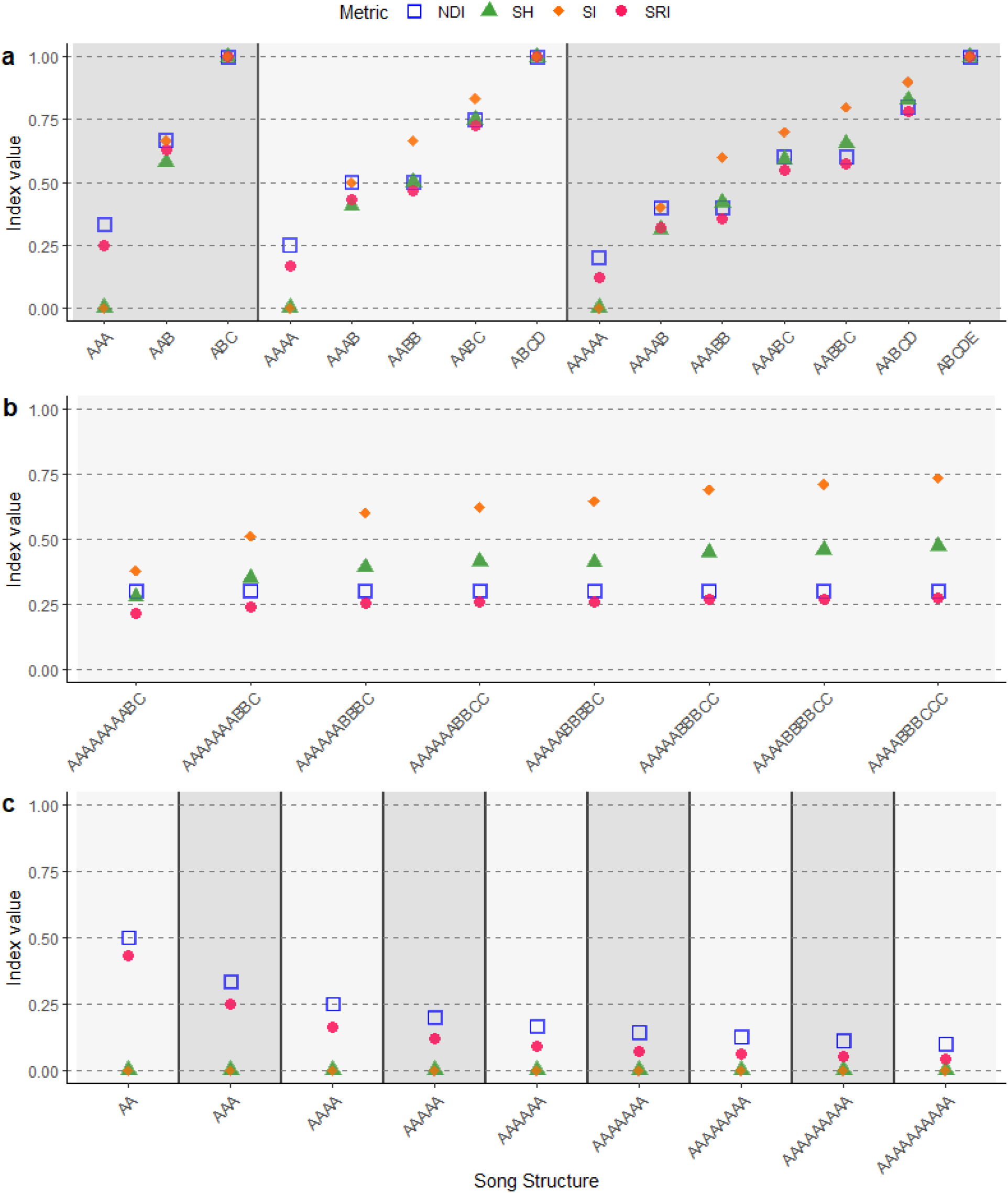
Comparison of three diversity indices using simulated sequences of songs: NDI-Note Diversity Index, SH-Shannon’s Equitability, SI-Simpson’s Diversity, and SRI-Song Richness Index. a) All structural combinations of lengths three, four, and five notes. b) Different length songs with same not repeating. c) All structural combinations of songs with ten notes and three note types. Each colored section represents a particular length of songs.

Similar patterns were observed for songs made of ten notes and three note types (eight structural groups of all possible combinations). For even repetition of notes, SH, SI, and SRI gave relatively higher values; but on the other hand, NDI provided the exact same value for all the songs (Fig. 1b)

Now, for all the songs made of a single note type but with varying numbers of notes, SH and SI provided the exact same value of zero. NDI and SRI could differentiate between these songs with the diversity values decreasing with increasing notes (more repetition) (Fig. 1c).

## Discussion

### Comparison with Note Diversity Index

The Note Diversity Index (NDI) provides non-zero diversity values to the songs with only a single note type repeating (Fig. 1c). But it fails to distinguish between the songs with the same number of notes and note types with varying repetitions (Fig. 1b). Song Richness Index (SRI) overcomes the limitations of NDI in these cases and gives different diversity values based on the number of times each note type is repeated.

### Comparison with Shannon’s Equitability and Simpson’s Diversity

Both Shannon’s Equitability (SH) and Simpson’s Diversity (SI) can distinguish between the various song structures with the same number of notes and note types (Fig.1b). But for songs with the same note repeating multiple times, SH and SI provide the same value-zero for the diversity (Fig. 1c). Given that in the case of any song, a bird puts a certain amount of effort in terms of notes which means all such songs cannot give the same value. SRI overcomes this limitation by providing a non-zero diversity value based on how many times the note is repeated and at the same time also able to distinguish between various song structures with the same number of notes and note types (Fig. 1a).

SRI can be an effective tool in quantifying the diversity of notes given its advantages on the traditionally used measures and no relative drawbacks. It estimates the diversity based on two main principles-i) more note types mean more diversity, and ii) more repetition of same note types means less diversity. SRI can have broader applications outside the acoustic analysis to calculate the diversity of combinations or sets with different elements. As it gives similar results as SH and SI with case-specific advantages, SRI can be used for any analysis that requires the measure of diversity.

## Acknowledgment

We thank Ishapathik Das of the Indian Institute of Technology (IIT) Tirupati and Anilatmaja Aryasomayajula of the Indian Institute of Science Education and Research (IISER) Tirupati for their constructive feedback while developing the new song diversity method. We thank the members of the two Ecology Labs at IISER Tirupati for their input and support. We acknowledge the support of IISER Tirupati.

## Data Availability

We provide sample datasets and the R code to calculate the Song Richness Index (SRI) in GitHub and Zenodo repositories. https://github.com/suyash-sawant/Birdsong-Diversity

